# CanDIG: Secure Federated Genomic Queries and Analyses Across Jurisdictions

**DOI:** 10.1101/2021.03.30.434101

**Authors:** L. Jonathan Dursi, Zoltan Bozoky, Richard de Borja, Jimmy Li, David Bujold, Adam Lipski, Shaikh Farhan Rashid, Amanjeev Sethi, Neelam Memon, Dashaylan Naidoo, Felipe Coral-Sasso, Matthew Wong, P-O Quirion, Zhibin Lu, Samarth Agarwal, Kat Pavlov, Andrew Ponomarev, Mia Husic, Krista Pace, Samantha L. Palmer, Stephanie A. Grover, Sevan Hakgor, Lillian L. Siu, David Malkin, Carl Virtanen, Trevor J. Pugh, Pierre-Étienne Jacques, Yann Joly, Steven J. M. Jones, Guillaume Bourque, Michael Brudno

**Affiliations:** DATA Team, University Health Network, Toronto, ON, Canada; Canada’s Michael Smith Genome Sciences Centre, BC Cancer Research Institute, Provincial Health Services Authority, Vancouver, BC, Canada; Princess Margaret Cancer Centre, University Health Network, Toronto, ON, Canada; McGill University, Montreal, Québec, Canada; Canadian Centre for Computational Genomics, Montréal, QC, Canada; Centre for Computational Medicine, Hospital for Sick Children, Toronto, ON, Canada; Hospital for Sick Children, University of Toronto, Toronto, Ontario, Canada; Division of Haematology/Oncology, The Hospital for Sick Children, Department of Pediatrics, University of Toronto, Toronto, ON, Canada; University Health Network, Toronto, ON, Canada; Department of Medical Biophysics, University of Toronto, Toronto, ON, Canada; Ontario Institute of Cancer Research, Toronto, ON, Canada; Département de biologie, Université de Sherbrooke, Sherbrooke, QC, Canada; Centre of Genomics and Policy, Department of Human Genetics, McGill University, Montreal, QC, Canada; Department of Human Genetics, McGill University, Montreal, Québec, Canada; Department of Computer Science, University of Toronto, Toronto, ON, Canada; Digital Products, Providence Health Care, Vancouver, BC, Canada; Zymeworks, Vancouver, BC, Canada; Sunbay B.V. Amsterdam, North Holland, Netherlands

## Abstract

Rapid expansions of bioinformatics and computational biology have broadened the collection and use of -omics data including genomic, transcriptomic, methylomic and a myriad of other health data types, in the clinic and the laboratory. Both clinical and research uses of such data require co-analysis with large datasets, for which participant privacy and the need for data custodian controls must remain paramount. This is particularly challenging in multi-jurisdictional settings, such as Canada, where health privacy and security requirements are often heterogeneous. Data federation presents a solution to this, allowing for integration and analysis of large datasets from various sites while abiding by local policies.

The Canadian Distributed Infrastructure for Genomics platform (CanDIG) enables federated querying and analysis of -omics and health data while keeping that data local and under local control. It builds upon existing infrastructures to connect five health and research institutions across Canada, relies heavily on standards and tooling brought together by the Global Alliance for Genomics and Health (GA4GH), implements a clear division of responsibilities among its participants and adheres to international data sharing standards. Participating researchers and clinicians can therefore contribute to and quickly access a critical mass of -omics data across a national network in a manner that takes into account the multi-jurisdictional nature of our privacy and security policies. Through this, CanDIG gives medical and research communities the tools needed to use and analyze the ever-growing amount of -omics data available to them in order to improve our understanding and treatment of various conditions and diseases. CanDIG is being used to make genomic and phenotypic data available for querying across Canada as part of data sharing for five leading pan-Canadian projects including the Terry Fox Comprehensive Cancer Care Centre Consortium Network (TF4CN) and Terry Fox PRecision Oncology For Young peopLE (PROFYLE), and making data from provincial projects such as POG (Personalized Onco- Genomics) more widely available.

## Introduction

With recent advances in bioinformatics and computational biology, there has been a massive expansion in the generation and use of -omics data such as DNAseq, RNAseq, methylomics and phenotypic profiling for research and in the clinic (Birney, 2019). With clinical sequencing becoming common practice, it is estimated that over 60 million patients will have their genomes sequenced in a healthcare context by 2025 (Birney et al., 2017). Analysis of -omics information benefits from large datasets yet handling the growing volume (Birney et al., 2017) of such data poses technical, security and privacy concerns, many of which are associated with the possible risk of patients’ re-identification. This risk must be balanced with the benefit that can come from the use of all data donated by research participants to promote scientific progress, and, possibly, improve healthcare (Knoppers and Thorogood, 2017). Thus, data custodians have the challenging responsibility to protect the privacy of patients and the security of their data, while maximizing data sharing.

While one of the most common ways to approach data sharing is aggregation of large datasets in central repositories, many national health frameworks, including those in Canada [4] and Switzerland, are inherently multi-jurisdictional. Within these countries, data from a single national study may have heterogeneous privacy and security requirements, with datasets of relevance having limited ability to move across jurisdictional boundaries to central computing systems. This makes accessing health data by researchers and clinicians challenging and time-consuming, as coordination across jurisdictions must be carefully done.

Data federation^1^ is being globally recognized as a solution to this challenge (Hermann, 2019), with queries and analyses “visiting” the distributed dataset while the items of data remain under local custodian control. Initial pushes towards interoperability and federation are driving new efforts and solutions (Birney et al., 2017). Internationally, the Global Alliance for Genomics and Health (GA4GH; https://www.ga4gh.org) has convened a number of national and international projects to share best practices and develop standards for interoperable data models and application programming interfaces (APIs). Several kinds of health data federations are active today and can be considered along three dimensions: how centralized/decentralized they are, the level of access they provide to the data, and the diversity of datasets accessible via the federation.

These three dimensions ultimately shape the way that data is stored and accessed within a given health data federation. First, the degree of decentralization (Figure 1) describes how queries flow through the system, and whether there are centralized or distributed identities. This includes “hub and spokes” models with a central infrastructure and identities, such as that of the Datashield project and the Local EGA Project (Fernández-Orth et al., 2019), and more decentralized, low trust models such as that of the Swiss Personalized Health Network (SPHN) (Froelicher et al., 2019, 2020; Raisaro et al., 2018). Second, data access can be approached in a variety of ways, from having a few predefined queries, such as with the Beacon Network [9] and the Matchmaker Exchange (MME) [10], to running arbitrary workflows, such as with Datashield [11,12]. This, in practice, depends on the governance of the federation, and determines authorization granularity and how results are returned. While authorization is commonly all-or-nothing at the level of datasets, federations which support deep access to data require finer-grained and more complicated authorization. In Datashield, for example, sophisticated approaches determine the sensitivity of fields and authorization can be made based on those calculations. Third, the federation can be designed for a particular data type (such as the original Beacon, which was a proof-of-concept considering only variants) or it can cover richer breadths of data types, ranging from multi-omics to clinical and phenotypic data to imaging. This is more valuable to researchers, but greatly increases the complexity, may include more sensitive data, and makes starting up such a federation a larger task. A careful balance between these dimensions must be struck in order for a data federation platform to be successful in a multi-jurisdictional setting.

**Figure 1:**
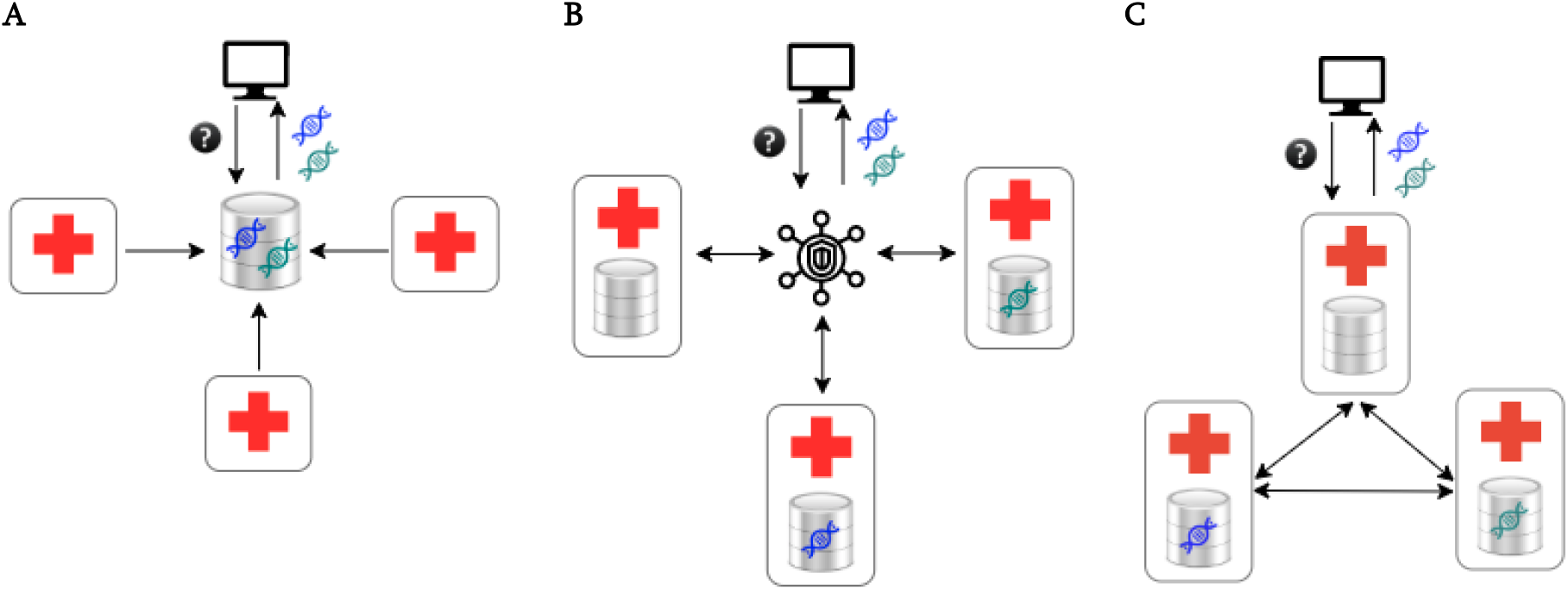
Degree of decentralization of a federated data platform. **A**: A centralized (not federated) data repository where data is pulled manually or automatically into a central data store. **B**: A hub-and-spokes model of federation, where there is significant central infrastructure that the peers are required to interact with. **C**: A platform that is a decentralized, peer-to-peer network, where a user sends a request to a peer health research centre - where relevant data may or may not be - and other peers are queried. Results (represented by DNA) are then returned to the user. If there were no connections between data sites and the user manually collected data from each site, we’d refer to this as merely distributed rather than federated.

Here we present CanDIG: the Canadian Distributed Infrastructure for Genomics. CanDIG is a national genomics and health data platform that carefully considers decentralization, data access and data types, as well as multi-jurisdictional privacy and security requirements to allow its users access to genomic and health datasets in any jurisdiction while maintaining local control of that data. It does this by implementing a unique federated approach that leverages existing technical infrastructure, GA4GH standards and clear division of responsibilities for its participants. Fast and straightforward data access and visualization are made possible with a flexible distributed dashboard that also allows for navigation of large genomic datasets and exploration of individual cases. Fine-grained security protocols determine the degree to which users can access data and enable authorized individuals to perform complex private queries for in-depth analysis using specific data of interest. Through this, CanDIG gives clinicians and researchers across the country secure access to the critical mass of data needed to understand and contextualize the myriad of -omics and health information being generated today. Built by a distributed development team across four institutions, and now deployed at five universities, hospitals, and research institutes across the country, CanDIG is hosting data for four national genomics projects, including data for 1053 patients with a variety of diseases as well as COVID-19 (see Table 1).

**Table 1:**
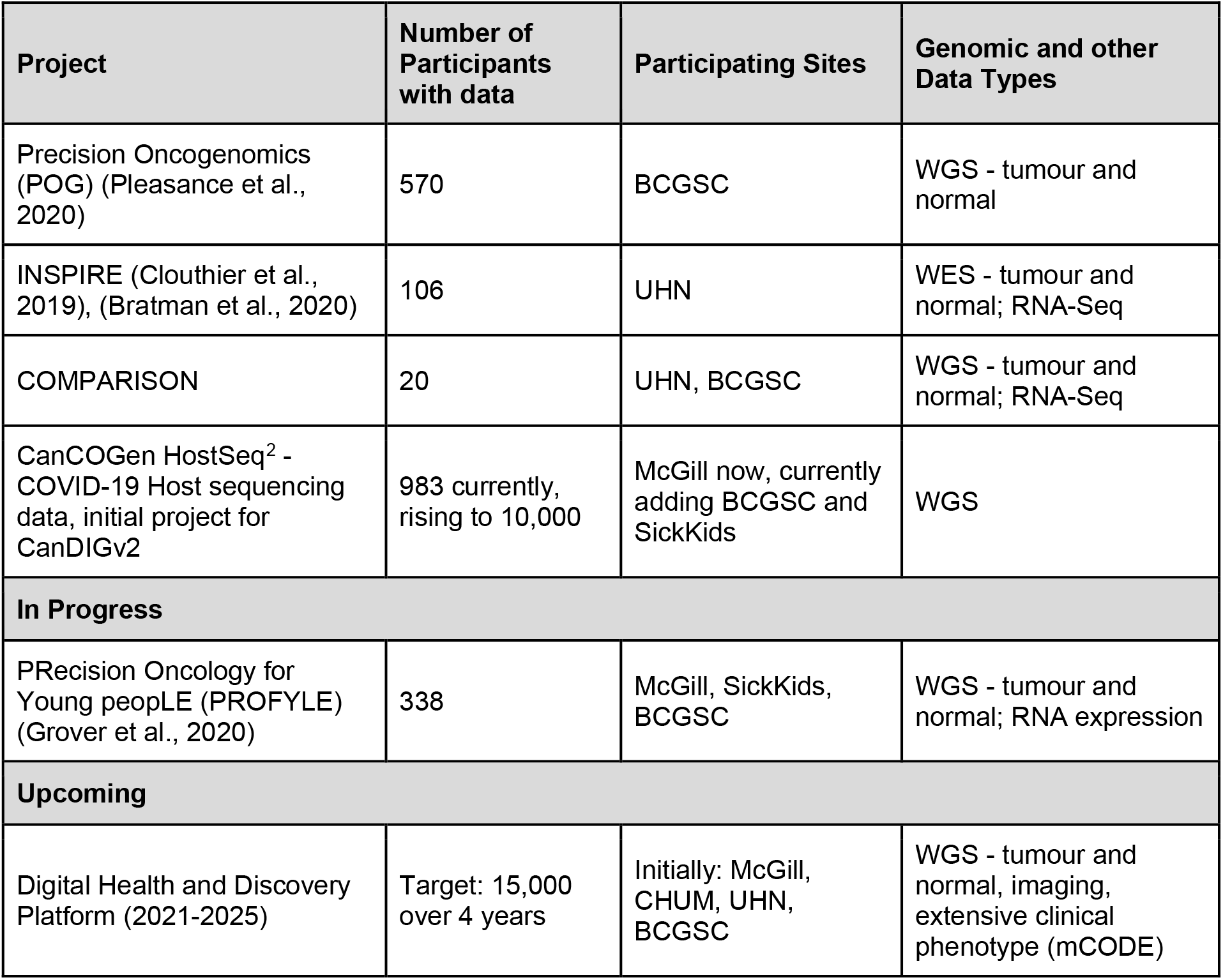
Current and forthcoming projects supported by CanDIG, including available patient numbers and data types. CanDIG’s federation and distributed authentication and authorization will be part of the DHDP^3^, a national platform initially supporting the government of Canada and Terry Fox Research Institute-supported Marathon of Hope Cancer Centre Network^4^, a 15,000-patient cohort-of-cohorts of consented cancer data.

## Results & Methods

### CanDIG Platform Overview

CanDIG is a fully distributed platform for human genomics and health data projects, and connects leading health institutions in Canada, including McGill University, The Hospital for Sick Children, University Health Network, Ontario Institute for Cancer Research, Canada’s Michael Smith Genome Sciences Centre, Jewish General Hospital and Université de Sherbrooke. CanDIG facilitates national research projects by building on existing technological infrastructure provided by Compute Canada, and by implementing a governance framework that divides responsibilities among its participants to provide secure data and metadata access. This enables institutional and provincial data custodians to judiciously allow discovery and analysis of data under their control by a national network of authenticated health researchers. As a GA4GH driver project, CanDIG is not simply a larger data silo; it uses international GA4GH standards, participates in their development, and interoperates with international partners as in the CINECA^5^ project, further expanding the pool of data potentially available to participants.

Each CanDIG site hosts the data and runs a local instance of the CanDIG software stack, which can be accessed by users at that site. Through this software, data custodians can make their datasets available to researchers at any CanDIG site for query and analysis, while still controlling and monitoring their access. This allows researchers to examine large, distributed pools of data in a uniform way.

A significant factor in CanDIG’s success at improving and streamlining data access for researchers is the implementation of clear divisions of responsibility between the platform, data custodians and data sites (Figure 2). The platform is responsible for best-practices software development for health data, interoperability with international projects, establishing governance and operations policies platform-wide, and onboarding data custodians. Data custodians are responsible for obtaining patient consents, data quality, removal of direct identifiers, and communicating authorization decisions from their data access committees to the platform. Data sites are responsible for following operations and security policy, keeping up to date with software releases, and contributing to the development effort.

**Figure 2:**
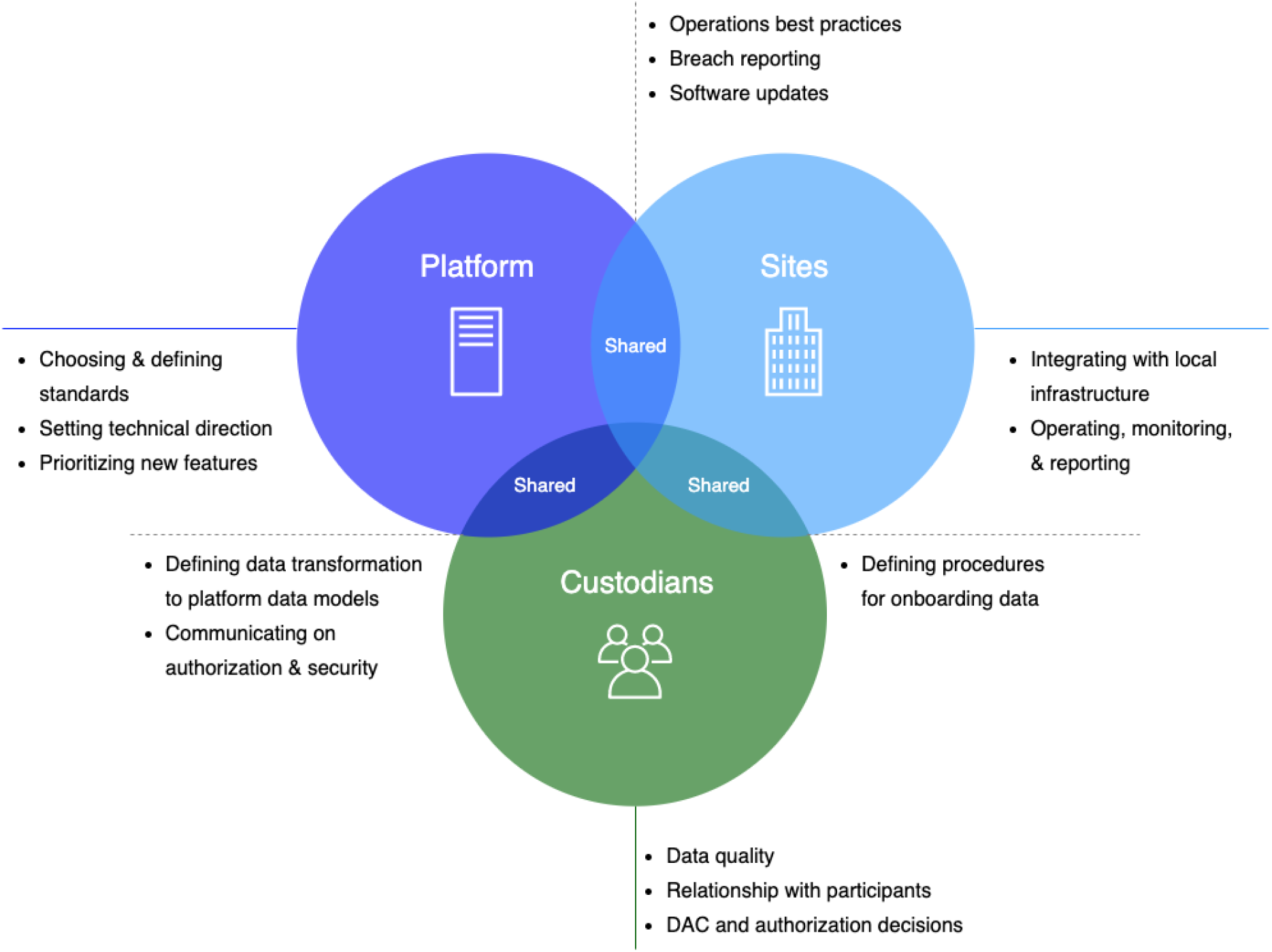
A high-level Venn diagram of roles and responsibilities of CanDIG Platform Partners - the data custodians (such as research projects), the platform effort, and the individual sites. A more detailed list of the responsibilities of each partner is shown in Supplementary Table 2.

CanDIG uses a governance approach that is flexible yet clear, specifying well-defined benefits and responsibilities for its stakeholders. Information about platform decision making can be found in the supplementary materials. It supports research on genomics data by enabling a wide range of queries and analysis services, and is completely distributed, with no central or coordinating infrastructure other than protocols, policies and governance agreements. It makes use of existing Compute Canada infrastructure, and gives CanDIG site operators complete and fine-grained control of access to all data entrusted to them, informed by platform-wide information services such as data access committee (DAC) authorization lists. Finally, to support the needs of data custodians, CanDIG makes health data available for analysis without directly sharing significant amounts of private data, even with other trusted sites. It also enables the use of differentially private aggregation queries for particularly sensitive datasets and allows granular data access audit capabilities. It supports project dashboards, interactive queries, and, increasingly, batchmode analyses.

### Use of GA4GH Standards and Technologies

By adopting GA4GH standards, CanDIG allows for easy startup of national and international multisite-omics projects where federated data queries and analysis can be made in a security-conscious environment. It makes genomics data available for querying and analyses through standard data models and modern ReSTful APIs. Making use of and participating in GA4GH standards development ensures that the solutions we build will be interoperable internationally, and allows us to take advantage of lessons, development efforts, and sometimes even entire components developed elsewhere. The current production CanDIG platform, CanDIGv1, implements GA4GH technical standards such as DUO^6^, RNAGet^7^, CRAM/SAM/BAM, and VCF, with CanDIGv2, in development, incorporating DRS^8^, WES^9^, service-registry^10^, Phenopackets^11^, Beacon (Fiume et al., 2019), and htsget (Kelleher et al., 2019), and shortly GA4GH Variant Representation (Wagner et al., 2021). How those technical standards fit together is shown in Figure 3. CanDIG is also adopting GA4GH standards such as the framework for responsible data sharing, data security and privacy policy, as well as security infrastructure recommendations. Implementation of these standards ensures that the datasets available through CanDIG can in theory be used by as many researchers, nationally and internationally, as possible, improving global study and understanding of health and disease.

**Figure 3:**
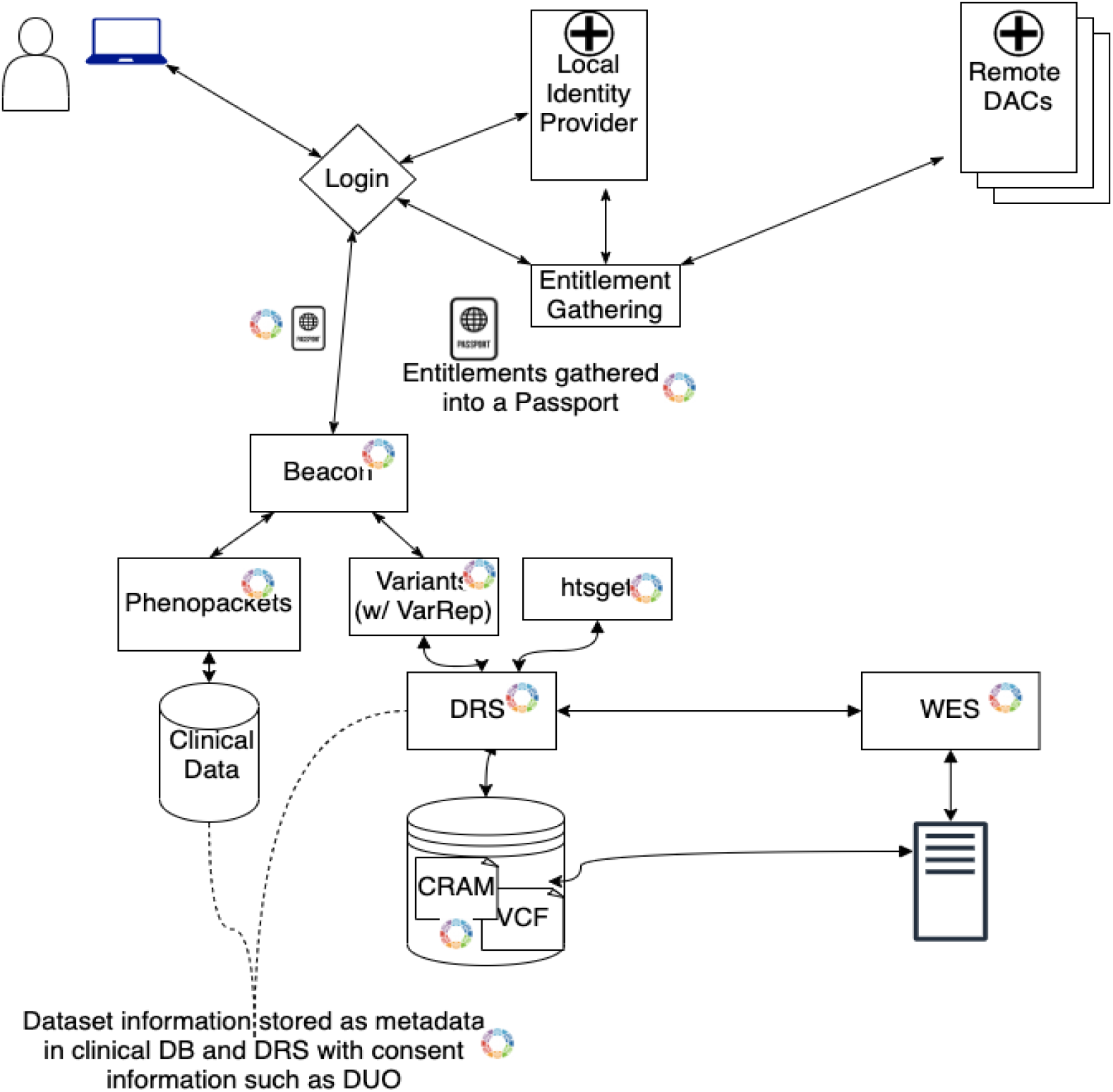
A schematic diagram illustrating how GA4GH technical standards and being used in the evolving CanDIG platform as we move towards deploying CanDIGv2. Illustrated are representations of services and standards being deployed within an individual CanDIG site, and how they build upon each other - use of standard Phenopackets eases the lightning of Beacons, a standard Variant Representation will avoid ambiguity of variants, the Data Repository Service Powers Workflow Execution Service as well as other file-based services such as htsget, and internally entitlements are propagated using a representation based on Passport visas. Consent metadata uses ontologies like DUO. Not shown are services following similar patterns as others, such as RNAget or refget, and that each service identifies itself using Service-Info. Also not shown are the use of policy standards such as the Framework for Responsible Data Sharing, Breach Reporting, or Privacy and Security Policy.

CanDIG’s standards-based approach and use of modern tools implemented elsewhere extends to infrastructure authentication, where we use OAuth2 and OpenID Connect to ensure the privacy and security of all data, relying on well-known implementations of Keycloak^12^ and Tyk^13^. By making use of trusted tools developed elsewhere as much as possible, CanDIG can focus on its distinguishing requirements of a decentralized federation. CanDIGv1 was originally based on the University of California Genomics APIs^14^, developed for an earlier version of the GA4GH effort, which allowed the CanDIG team to immediately make progress on federation.

### CanDIG’s data federation design

CanDIG’s choices of data federation design harness policy and technical infrastructure already existing in Canada and elsewhere. That includes having a modest number of trusted institutions as participants, who have a history of working together, each with their existing well-established hardware, operations, and data governance, as well as several active research projects, which are often national. Leveraging pre-existing infrastructure and research networks made initial implementation of CanDIG efficient, and in the future will allow for easier integration of new participant sites. We provide a comparison of the CanDIG design with that of other data federations, along the three dimensions of decentralization, data access, and range of data types in Figure 4. (When designing our approach from the ground up, we also find a more granular six-dimension approach to be useful, which is described in Supplementary Table 1.)

**Figure 4:**
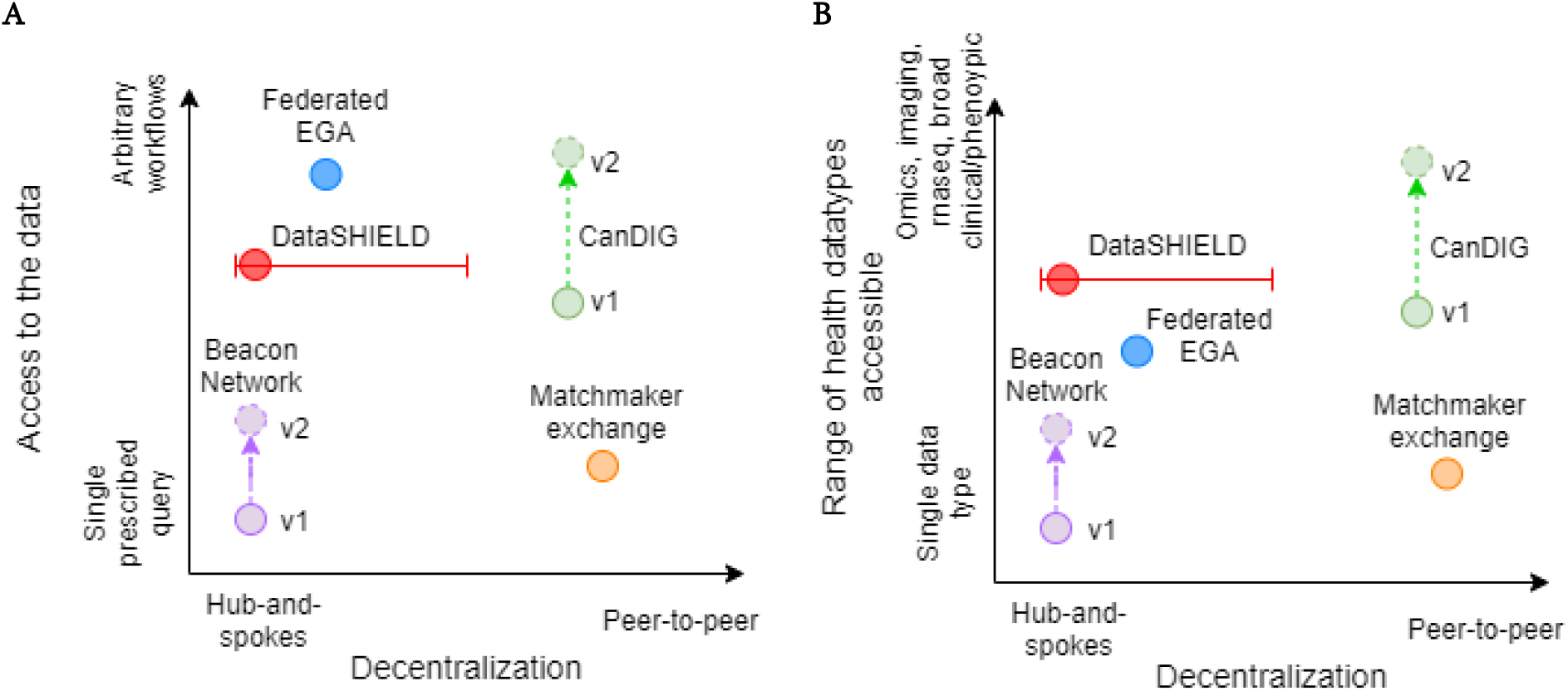
Categorizing well-known health data federations along dimensions of decentralization (hub-and-spokes to peer-to-peer) and, in the left panel, level of access to data (pre-specified queries to arbitrary workflows). In the right panel, the range of different kinds of datasets supported.

Heterogeneous health data privacy requirements means that central “hub” infrastructure used by some federations was not desirable for the development of CanDIG and pushed us towards a fully distributed approach. While a hub-and-spokes approach can be technically easier in such a situation, in our context it would introduce a number of governance concerns, such as who owns and is responsible for the central infrastructure, how central identities are to be handled, and which jurisdiction’s set of privacy requirements applies to the central hub. The distributed nature means we can use nationally-recognized local identities, with local authorization decision-making and data-gathering delegated to institutions.

With respect to data access, taking advantage of existing agreements and infrastructure while remaining fully distributed means that we benefit from being able to rely on a strong high-trust closed governance model. This facilitates quite deep “visiting” of the local data stores, if locally allowed, pushing us towards rich APIs that support such ad-hoc analyses (Figure 4).

Improved understanding of health and disease requires contextualization of patient data through multi-omics approaches and research initiatives that utilize a broad array of data types. To facilitate the wide range of research projects and queries we want to enable pushed us to support an increasingly rich range of datasets. This, along with the range of data access that CanDIG enables, requires granular control over access to data and the development of services which can support multiple data types.

Our design choices, with respect to decentralization, degree of data access and the types of data available, allow CanDIG to support genomics research needs of research project managers, researchers, and clinicians in a multijurisdictional environment through powerful project dashboards, browsers for variants and genomic data, and complex queries across our federated datasets. These are described in turn, below.

### CanDIG Project Dashboard

The CanDIG dashboard and APIs have to support a number of different users and their use case. Perhaps simplest is that of the research project manager, monitoring data as it is being collected onto the CanDIG sites, examining summary statistics, and checking quality of input data.

Simple aggregations like count of patients by geography, phenotypes, and data types available can be accessed at low levels of authorization for a dataset via APIs or visualized directly through a project dashboard. Dashboards pages show simple status counts, breakdowns by category, and overviews of data sets (Figure 5A-C).

In our decentralized federation, there is no central “CanDIG” identity; each user has a home site, typically the institution at which they work, where they can log in using their local credentials, and can view the dashboard there. This propagates the queries necessary to drive the dashboard across to all federation partners (Figure 1A). Currently a simple, single-step fan-out is sufficient. Each site in the federation recognizes the identity of the other sites and makes authorization decisions regarding access to the data it hosts. The granularity of that authorization allows for multiple projects to be supported without exposing data from one project to researchers of another unless so authorized. That information is then presented to the user via their home site. Because the user is necessarily authorized to see the data from the response, and the home site is effectively the user’s work computer, the requests can be safely aggregated at the home sites.

**Figure 5:**
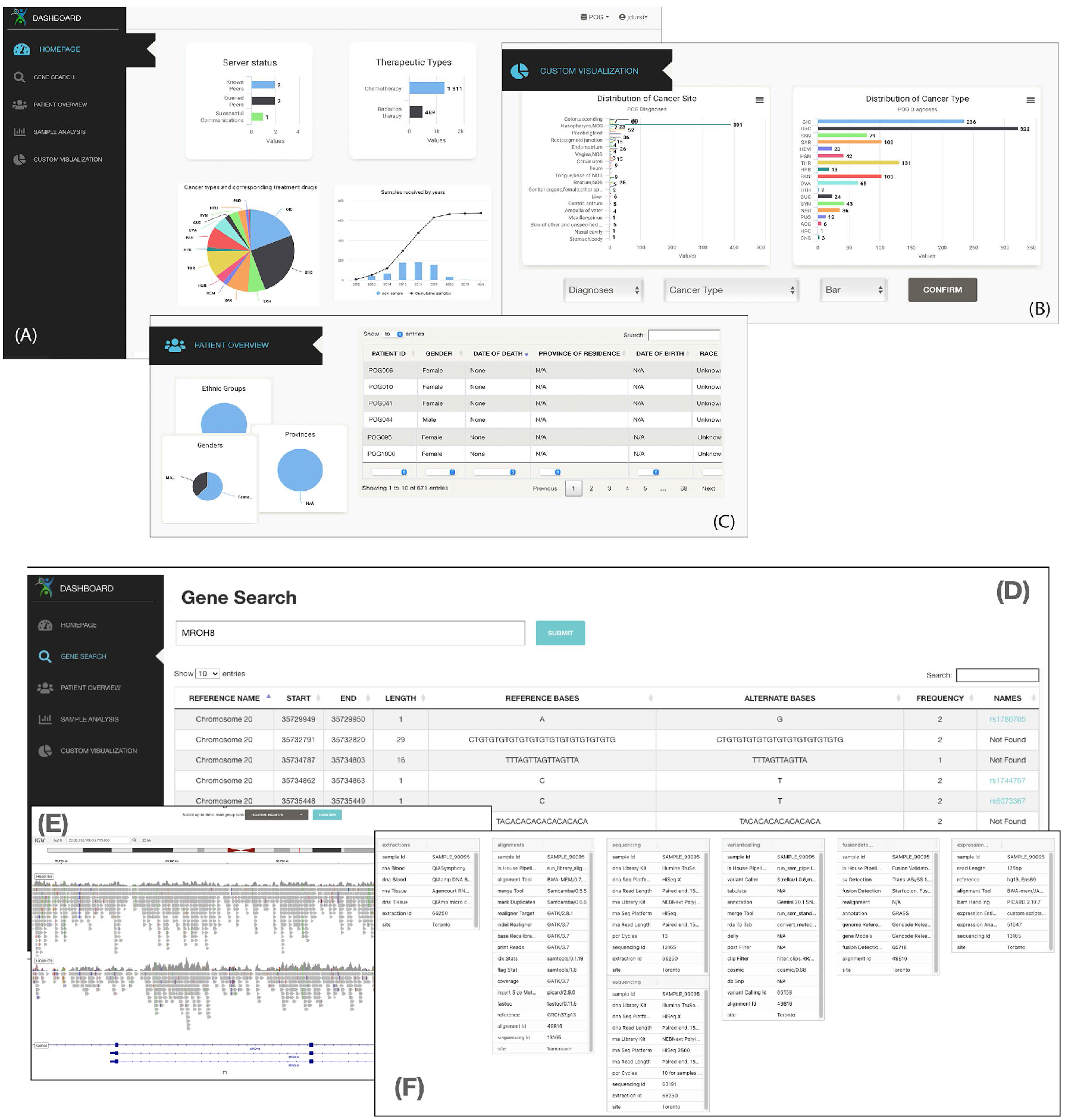
CanDIG dashboard overview pages for a research project manager, or someone examining a dataset for the first time - shown is the dashboard for the POG project, with credentials that only allow limited data access. The website, and in particular the dashboard, is reactive and mobile-friendly, so that project status can be checked from anywhere. In panel **A**, the default simple summary statistics about participant data and cases available on the platform is shown. A project manager who wants to see breakdowns by other variables can use the custom visualization widget, illustrated in panel **B**. Finally, the research project manager can quality check clinical data by patient or check to see what associated genomic data is in the system through the patient’s tab, illustrated in panel **C**. A bioinformatician working with a particular sequencing project will want to explore genomic variants and read-level data, shown here with a federated mock dataset. A gene search page, shown in panel **D**, allows querying variants in the dataset by gene name; from there the bioinformatician can view reads from the variant location in one or more participant’s data in an embedded IGV.js tool, shown in panel **E**. Information about the analysis pipeline used to generate the data from the sample collection to the variant calling tool settings can be shown through the sample analysis tab, shown in panel **F**.

While this is our current approach, CanDIG is an ever-evolving platform, and other topologies have been tested and can be used as the number of our participating sites grows.

### Browsing Variants and Genomic Data

Finally, researchers who have to verify data quality can have very deep access to the data, allowing them to dig into individual cases. This can be via API access, or via the dashboard which allows viewing of mutations by gene (Figure 5D), integrated IGV (Robinson et al., 2011) for viewing variants and their sequencing context (Figure 5E), and information about the analysis pipeline producing those results (Figure 5F).

### Enabling Complex Private Queries across CanDIG

The CanDIG platform provides authorized researchers access to complex queries and analyses of distributed datasets, jointly across various supported data types. This is enabled through very fine-grained authorization. Staff and researchers with different project roles or areas of specialty are granted different “access levels” to a dataset by the study’s DAC, with each record of clinical and phenotypic data belonging to a dataset and each property within the record having a default access level. However, the required access levels of each item - each property of any particular row - can be increased, allowing, for instance, data custodians responsible for data from particularly marginalized populations to require additional levels of authorization to access the study data.

With the authorization in place, complex queries that researchers wish to perform programmatically or that are awkward to use a web interface for are available through increasingly rich APIs connecting the individual services and APIs supported by the platform. These components were unified through a single `/search` endpoint returning the information, or `/count` which returns counting aggregations.

The queries serve both discovery and simple analyses use cases. As an example of a discovery use case, a user can search across datasets to find patients with a given phenotype, genomic variant, and clinical information, such as adult cancer patients with TP53 mutations who have undergone treatment with platinum-based therapies. If a clinician-researcher is authorized to access the data, this allows them to dive directly into a particular case (as shown in the previous two use cases). For custodians who wish to make data available for discovery queries to a broader range of researchers while still maintaining participant privacy, CanDIG supports these queries as counts with local differential privacy (Duchi et al., 2013). Laplace or exponential noise is added to counts at each site before being summed and presented to the user (Figure 6A). This is the “local” addition of differentially private noise (as opposed to global differential privacy, Figure 6B), it allows the custodians of data at each site to determine the privacy-utility tradeoffs for the data under their control. The results of these discovery queries can then be used to guide subsequent data access requests for higher levels of authorization to a dataset, or for secondary use access to a subset of the data. CanDIG discovery queries are available in other environments such as through the GenAP platform for this purpose.

**Figure 6:**
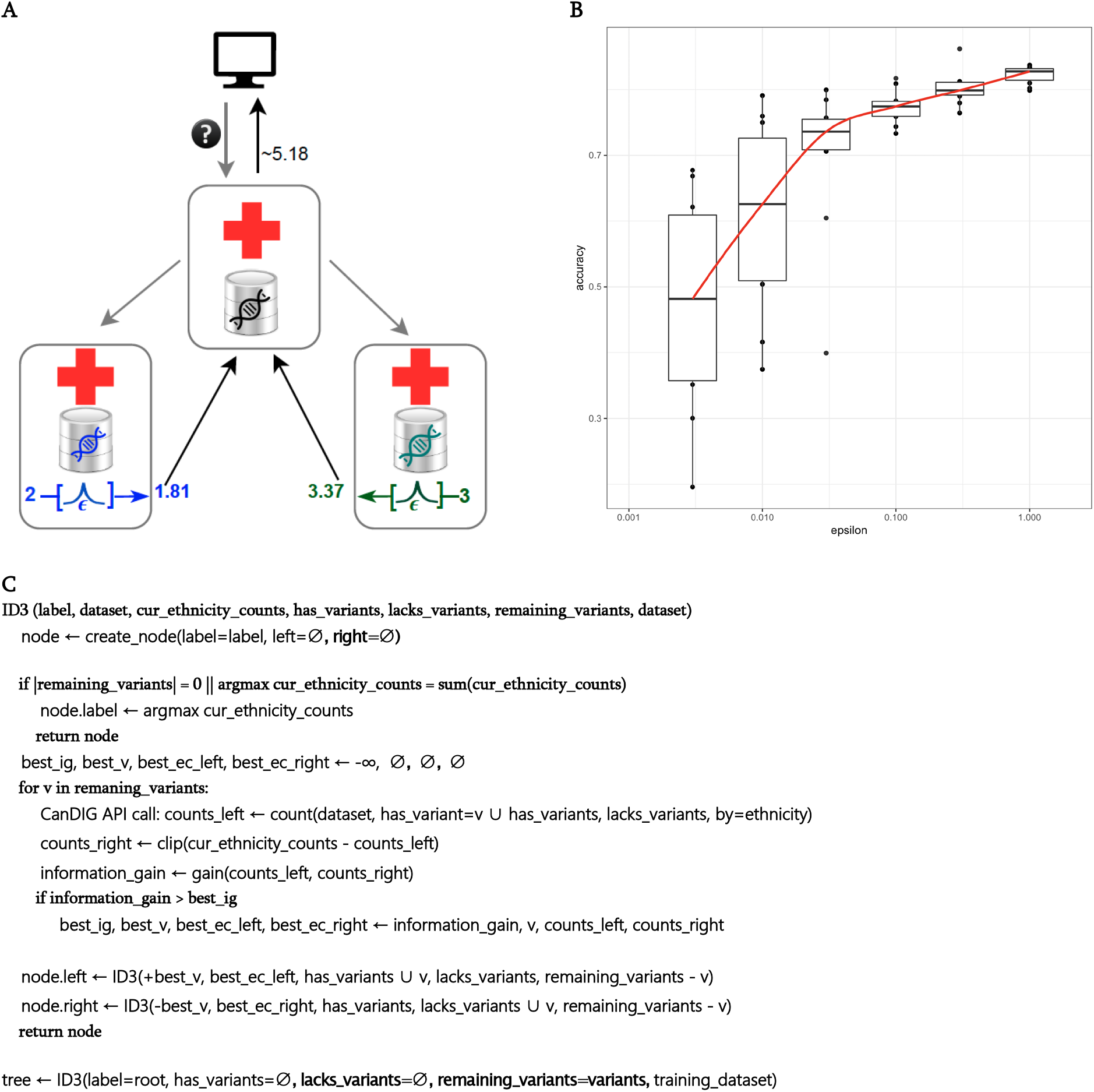
**A** illustrates the implementation of differential privacy in the CanDIG federation. Consistent with our approach of ensuring local control over authorization decisions, for queries or datasets where differential privacy is required, we implement local differential privacy. In local differential privacy, each data site determines its privacy parameter e and adds the noise. Lower e means lower privacy loss, requiring more noise and lower resulting accuracy. This contrasts with global differential privacy, where a central site adds a global level of noise after getting unperturbed data from the sites. **B** demonstrates a proof of concept, training of an ID3 tree based on 17 known ancestry informative SNPs on five the 1000 genomes superancestries, with per-query laplacian noise (determined by the query sensitivity and ***ϵ***) added at each query, trained on 250 randomly selected training genomes and tested against held out samples. With decreasing levels of the privacy parameter ***ϵ***, privacy increases and utility is diminished; accuracy varies more widely run to run with larger amounts of noise (smaller values of ***ϵ***). **C** shows pseudocode of the client-side ID3 algorithm, making count calls to the CanDIG APIs.

The APIs expressiveness is enough to support building more complex analyses on the client side. For instance, it is possible to train ID3 classifiers (Quinlan, 1986) to predict ancestral data from a modest number of informative SNPs in the 1000 genomes data using the existing counts functionality, with or without differential privacy^15^. Here we perform queries on the client side that gather counts of a small set of known-informative SNPs - a subset of 17 out of a known panel of 55 (Kidd et al., 2014), grouped by piece of demographic information (the 1000 genomes ancestry), and train a decision tree classifier based on the “splits” of ancestry by presence or absence of the SNPs (Figure 6C). ID3 decision trees work particularly well with differentially private aggregations due to their use of counts.

More simply, it is possible to use the APIs in a similar manner to perform more traditional population genetics visualizations on the 1000 genomes data such as identifying SNP counts versus ancestry. This work is informing the development of new APIs to allow such analyses to be done with less development by pushing the analyses to the server rather than the client. This will also serve to focus development of backend services to improve the performance of these utility machine learning (ML)-supporting queries. CanDIG v2, currently under development, supports the GA4GH WES 1.0 standard^16^ for workflows to run on datasets for authorized users.

## Discussion

Researchers and clinicians need appropriate tools to access and analyze rapidly-accumulating - omics data in an informative context. Such a context is best achieved through the use of large and comprehensive data sets, which allow for thorough investigations to shed light onto various clinical and research questions. Meanwhile, in distributed environments, especially those that are multi-jurisdictional, local and national project managers and data custodians need to ensure that their data remains securely and privately under the appropriate domain of control. As the first Canadian national health genomics data federation platform, CanDIG lets researchers and clinicians in Canada access and benefit from a national pool of -omics data and enable much simpler sharing of data across projects - both nationally and internationally. By connecting multiple national health data projects in Canada on a single platform compliant with GA4GH standards, CanDIG is enabling all to benefit from the data being generated.

The governance of our fully distributed, multi-jurisdiction, and multi-project platform is complex. While the roles and responsibilities of different overlapping stakeholders can stay implicit in simpler centralized or single-dataset projects, in CanDIG they are made explicit. There are significant advantages to this, despite requiring more work up-front for delineating the roles and responsibilities of the platform, the software development effort, the sites, and the data custodians. This clarity and separation of roles has greatly eased participation in international federation efforts such as with the EU/Canada/Africa CINECA project^17^. We believe that our approach is portable to a number of other distributed health data projects that share data among trusted partners.

Based on what we have learned, we are continuing to develop CanDIG with a more scalable service-oriented architecture with support for workflows, policy-driven authorization, data access committee portals such as REMS (Linden et al., 2013), more molecular data types, and more complex analyses supported by additional APIs and “map-reduce” type federation of queries. We are also looking to enable interoperability with clinical data by moving to a standard clinical data model (OMOP) (Overhage et al., 2012)), querying of medical imaging metadata, and stronger ontology support. These additional capabilities will be necessary to support the upcoming DHDP project, with greater scale, more collaboration, and more data types - technical papers describing this work are in preparation - but the fundamental distributed authentication, authorization, and federation approach will remain essentially unchanged. In addition, preparing for more sites joining the CanDIG federation, we are strengthening the project’s governance structure, expanding on our platform consortium governance policy, site-platform agreements, and platform-sequencing project agreements.

CanDIG allows for improved access and analysis of multi-omics data on a national level. As genomics becomes more affordable and accessible, we are able to use it to better inform clinical decisions and improve patient outcomes, particularly in cancer and rare disease. Through a national federated platform such as CanDIG, we are able to provide researchers and clinicians with higher quality data and more in-depth and complex analyses. This ultimately allows them to extract even more information from the datasets that are made available to them, thus further advancing diagnostics and patient care.

## Acknowledgements

CanDIG development was funded by the Canada Foundation for Innovation Cyber infrastructure grant 34860, CANARIE Research Data Management contracts RDM-090 (CHORD) and RDM2-053 (ClinDIG), and the Canadian Institutes for Health Research as part of the Africa-Canada-EU Horizon2020 CINECA project (CIHR grant number #404896). The CanDIG team would like to acknowledge the standards and convening work of the Global Alliance for Genomics and Health, and the work of the University of California at Santa Cruz’s Genomics Institute, particularly (but not exclusively) Danny Colligan, Jerome Kelleher, and David Steinberg, for the development of the ga4gh-server which formed the starting point of our prototyping. The CanDIG team would like to acknowledge the participation of the PROFYLE program made possible through funds from The Terry Fox Research Institute, Air Canada Foundation, Alberta Cancer Foundation, Alberta Children’s Hospital Foundation, BC Cancer Foundation, BC Children’s Hospital Foundation, CancerCare Manitoba Foundation, Charles-Bruneau Foundation, CHEO Foundation, ChildCan, Childhood Cancer Canada, CHU Sainte-Justine Foundation, Dalhousie Medical Research Foundation, Fight Like Mason Foundation, Garron Family Cancer Centre, IWK Foundation, Janeway Children’s Hospital Foundation, Kids Cancer Care Foundation, McMaster Children’s Hospital, The Montreal Children’s Hospital Foundation, Phoebe Rose Rocks Foundation, Sarah’s Fund for Cedars, SickKids Foundation, SickKids Research Institute, and Team Finn Foundation. GB is a Canada Research Chair in Computational Genomics and Medicine. MB is a CIFAR Canada AI Chair.

## Supplementary Materials

### Design Principles

Underlying CanDIG is a set of design principles which shape both project and technical decision making.

CanDIG is fully **distributed**: All data, and all infrastructure, is completely distributed; no shared or centralized services. All coordination is done at the level of policy, protocol, or software development.

CanDIG must support full **local control**: Consistent with common governance and policies, local data custodians have complete control over access to their data, and auditability/observability into data access and use. We do not support large-scale data transfers and downloads as they sacrifice later auditability.

CanDIG is **API based and driven**: Since we are building a platform whose success hinges on interaction between users and multiple sites, new development will rely on API-first design, with APIs developed and documented, and services and clients built on this. This ensures documentation of the APIs, interoperability between clients, and alignment with GA4GH efforts.

CanDIG emphasizes **security and privacy**: Our mission is to connect privacy-sensitive (although not directly identifying) human health data. We will rely on modern authentication and authorization technologies such as OpenID Connect so that secure API-based access to data can be achieved. We will use well-trusted tools wherever at all possible for security-sensitive code. We follow GA4GH Security Working Group’s best practices. Since CanDIG is also fully distributed there is no central infrastructure to maintain or secure. Methods such as differential privacy(Dwork, 2011) are baked into the project from the beginning.

**Open Source, Standards-Based, and Interoperable**: CanDIG builds on existing standards, on matters both technical (OIDC, OpenAPI, ReST, Docker, etc.) and genomic (via its role as a driver project for the GA4GH effort). This approach allows interoperability as wide as possible, while focussing efforts only where it matters most. Rather than building a single larger silo, we intend to <specific description of how CanDIG differs from data silos>.

#### CanDIGv1 Technical Prototype

Our work in v1 focused on developing the fundamental fully-distributed, API driven platform, codesigning and validating it with real health genomic sequencing projects. Given the scale of the initial deployment, any components which could rely on manual processes were implemented in that manner while the key data federation components were developed.

We took as a starting point the UCSC genomics API implementation, `ga4gh-server`^18^. This provided the initial API layer for building our fully distributed and API driven platform. From here we began implementing a wider range of APIs and data models to allow more complex queries, richer clinical and phenotypic data, and project dashboards for data custodians.

To support security and privacy we implemented stronger OIDC authentication, moving responsibility for the “OIDC Dance” to an authenticating reverse proxy, Tyk, which allowed us to begin the migration from the original monolithic application to a service-oriented architecture; this also enabled the integration of additional stand-alone services such as GA4GH standards Beacon (Fiume et al., 2019), htsget(Kelleher et al., 2019), and rnaget. We also implemented multiple differential privacy APIs to allow data custodians to make their data available to less highly-authorized researchers while maintaining strong privacy guarantees.

To fully support local control, our authorization focused initially on a local site maintained whitelist of researchers and their data entitlements (the data projects and access levels for their authorization within each project). In work for the next version of CanDIG we are implementing platform wide DAC authorization lists which will be added to (and possibly overridden by) the local authorization lists. The roles of the platform and site in authorization and data access are defined in a deepening set of governance documents for the project and the platform.

Sites have control not only over data authorizations but also the technical operations of their site; thus, we need clear service boundaries defining where the responsibilities of the CanDIG stack stop and those of the site’s operations team begin, and those boundaries need to be defined in a way that provides common interfaces to the CanDIG stack while providing maximum flexibility to the site. In v1, a key service boundary is in the site’s authentication services; we prescribe Keycloak (Christie et al., 2017) as a service for each site to provide OIDC tokens for its authenticated users, but how local identity management is handled is defined by the site’s infrastructure and needs.

The remaining service boundaries are straightforward in v1 but work underway for the next version which will support larger scale and automation, we have defined new boundaries. MinIO and GA4GH Data Registry Service (DRS) mark the standard APIs for storage which the CanDIG stack can leverage but allow sites to use a variety of back-end storage systems; Similarly, the GA4GH Workflow Execution Service (WES) [cite] and the Common Workflow Language (Amstutz et al., 2016) (CWL) allow us to decouple the definition and invocation of computational pipelines from how they are run on back-end computational infrastructure.

To support standards and interoperability, we build wherever possible on existing technical and genomic standards. As mentioned earlier we use OIDC for authentication, and OpenAPI for defining APIs; we rely on standard genomics formats like CRAM and VCF; we use and help shape GA4GH APIs such as Htsget, Beacon, and RNAget, GA4GH ontologies such as the Data Use Ontology^19^ (DUO) for consented use for data; and as mentioned before, immediately upcoming work will use standards in exposing services such as CWL, GA4GH WES, and DRS. We have established the success of this approach in promoting interoperability by demonstrating initial twoway authentication with the ELIXIR Authentication and Authorization Infrastructure (Linden et al., 2018).

### Data Federation in the CanDIG model

We have found useful a six-dimension approach to describing the design of data federations, using the above distinctions. Foundationally, a governance model which includes how federation peers join and leave, and the trust model required between them; how authentication of federation users work; the granularity of authorization; what queries are enabled; how queries flow through the federation; and how intermediate data is combined to be presented to the researcher.

**Supplementary Table 1:** a comparison of different federation models along six dimensions - governance/peer trust, authentication, authorization granularity, query range, query flow, data gathering.

| Model | Governance & Peer Trust Model | Auth'n | Auth'z granularity | Queries | Query flow | Results gathering |
| --- | --- | --- | --- | --- | --- | --- |
| Beacon Network (Fiume et al., 2019) | Open | Partial recognition of external identities | Multi-dataset; open, registered access, controlled access | Single-query | Hub-spoke | Aggregation to hub |
| MME (Philippakis et al., 2015) | Closed | Site-to-site | Single-dataset | Single-query | Pairwise | Pairwise aggregation |
| DataShield (Gaye et al., 2014) | Somewhat closed; peers trust central portal | Central identity | Fine-grained | Many (~140) primitives | Hub-spoke | Summary statistics only |
| Local EGA (Fernández-Orth et al., 2019) | Closed | Central identity | Multi-dataset | File access | Hub-spoke | Aggregation to requestor |
| SPHN (Raisaro et al., 2018) | Closed, low-trust | Recognition of node identities | Fine-grained, multi-dataset | Multiple APIs | Peer-to-peer | Secure multi-party computation, homomorphic encryption |
| CanDIG | Closed, high-trust | Recognition of peer identities | Fine-grained, multi-dataset | Multiple APIs | Peer-to-peer | Aggregation to requesting site; optional differential privacy |

In MME, researcher identities are only meaningful at each node, and the nodes themselves make requests of each other.

The depth of access to data often reflects deeper cooperation and shared governance between the data sites. Federations can be quite open, allowing new peers readily, such as the Beacon Network, or closed, allowing deeper access to the data or access to sensitive data but requiring formal agreements to be signed to join (such as Datashield, with deep access to data, or MME with access to deeply phenotyped childhood rare disease data).

We have a strong trust model between our participating institutions. They are all teaching hospitals and research institutions with long histories of collaboration and experience signing and honouring agreements with each other. A primary use case for CanDIG is to support national projects that the sites are collaborating on. While we do not expose raw data between sites, this strong level of trust gives us greater flexibility in the release of and combination of intermediate results in analyses.

Platform authentication is performed at the institution level - each institution provides a strong identity for its CanDIG users, and each user must have a CanDIG institution vouching for them. All requests in the platform are tied to a single user, ensuring enforceable accountability at the level of the researcher and institutions. This approach means that all users of the system follow the Registered Access (Dyke et al., 2016) access model. For authentication we use the web— standard OpenID Connect (Sakimura et al., 2014) (OIDC) technology.

For controlled access data, access authorization decisions are made locally by the data sites. In our case, these sites bear ultimate responsibility for incorrectly authorized data release. However, many of the data sets stored within CanDIG are part of larger national projects which have data access committee lists maintained by one of the sites. Thus, the local authorization takes as input external data.

Query flow through our system is entirely peer-to-peer. Since every user “belongs to” an authenticating site, their queries can flow to that site, whence they propagate outwards through the closed federation to be received at peer sites. All requests through the system are associated with the single authenticated researcher who made the request; this is simplified by our adoption of OIDC authentication. We have experimented with peer-to-peer cycle topologies to enable certain types of privacy-enhancing aggregations but for the time being we use a simple fan-out topology.

For result aggregation, we make use of the fact that each user is strongly associated with a particular site, and so results the user is entitled to access can be safely aggregated on that site. We have also experimented with the use of homomorphic encryption to allow for global approaches to differentially private aggregations. This requires lower amounts of total noise (and thus higher result utility) to maintain privacy than our current local (or regional) differential privacy approach but is not implemented currently.

Finally, CanDIG version one (v1) does not automate any form of platform-wide observability or auditability of access patterns across the closed federation as a whole, as given the small number of sites, the closely knit team, and modest query rate this has not been necessary. Approaches are being proposed for v2.

### Roles and Responsibilities in the CanDIG model

With a “multi-tenant” platform supporting multiple research projects, and federation separating the roles of sites and the platform as a whole, the CanDIG project has had to be very explicit about the roles and responsibilities of each partner in its role. Below in Supplementary Table 2 is a more detailed listing of the roles and responsibilities outlined in Figure 2.

**Supplementary Table 2:**
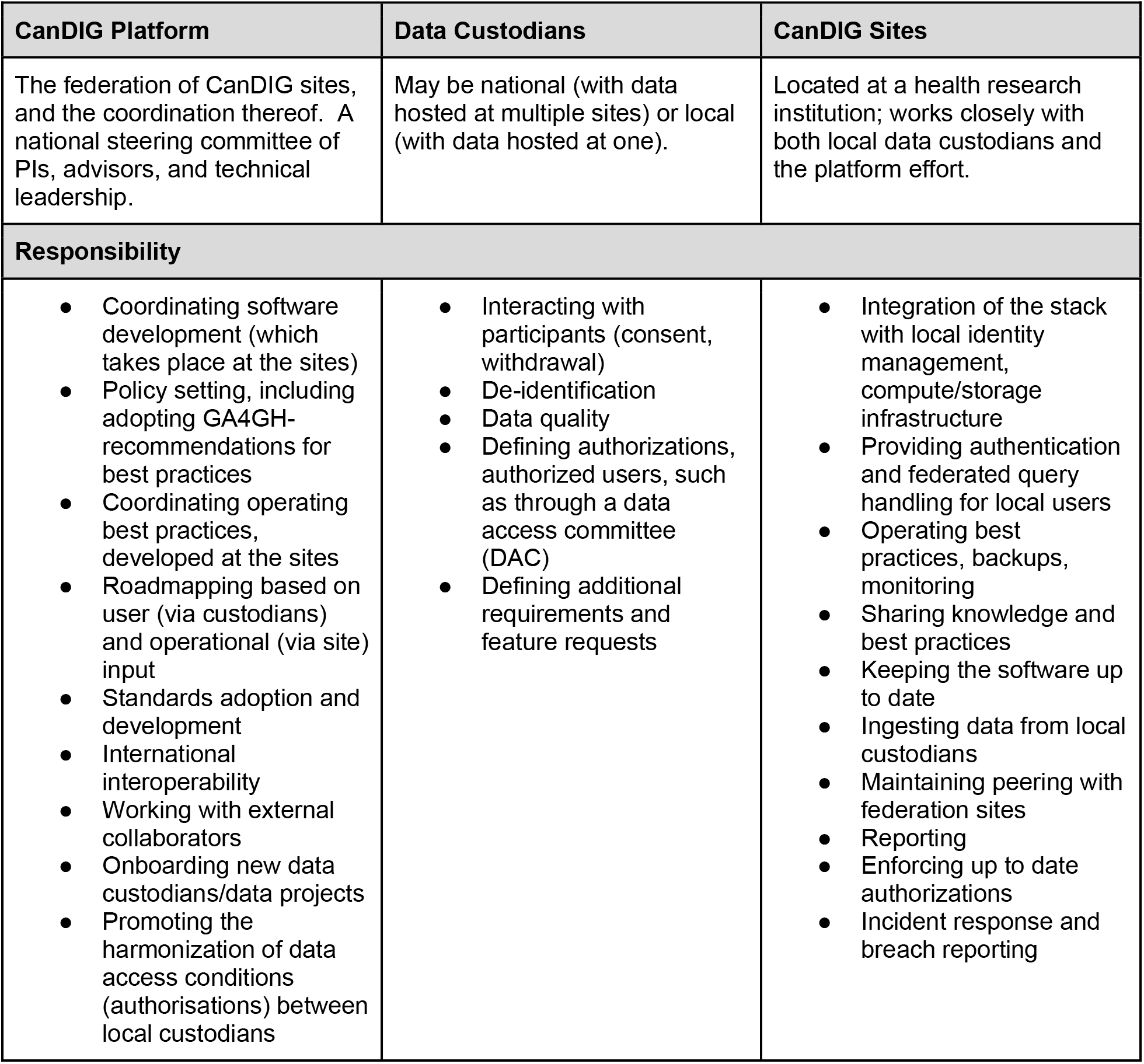
Detailed listing of roles and responsibilities of CanDIG partners.

## Footnotes

1 We mean something quite specific here by data federation - querying and analyzing horizontal partitions of data, with participants geographically separated and all of their data located at one site. We do not consider here linking multiple separate data sources for the same data subject (e.g., clinical data in one store, genomic data in a second store, crossing “vertical partitions”); in our model this happens internal to a site and we refer here to those operations as performing data integration rather than data federation.

2 https://www.genomecanada.ca/en/cancogen/cancogen-hostseq

3 https://www.dhdp.ca/

4 https://www.marathonofhopecancercentres.ca/

5 Common Infrastructure for National Cohorts in Europe, Canada, and Africa: https://www.cineca-project.eu/

6 https://github.com/EBISPOT/DUO

7 https://github.com/ga4gh-rnaseq/rnaget-compliance-suite

8 https://ga4gh.github.io/data-repository-service-schemas/

9 https://ga4gh.github.io/workflow-execution-service-schemas/

10 https://github.com/ga4gh-discovery/ga4gh-service-registry

11 http://phenopackets.org/

12 https://www.keycloak.org/

13 https://tyk.io/

14 https://github.com/ga4gh/ga4gh-server

15 https://github.com/CanDIG/ID3_Project

16 https://github.com/ga4gh/workflow-execution-service-schemas/releases/tag/1.0.0

17 http://www.cineca-project.eu

18 http://github.com/ga4gh/ga4gh-server

19 https://www.ebi.ac.uk/ols/ontologies/duo

